# UniMedSeg: Unified Multi-Modal Medical Image Segmentation via Uncertainty-Guided Encoding and Frequency-Spatial Dual-Prompt Decoding

**DOI:** 10.64898/2026.10.03.755900

**Authors:** Jingling Zhang, Shuting Zheng, Xiangfei Liu, Wen Zhang, Jia Gu

## Abstract

Text-driven medical image segmentation demands unified models based on multi-modal learning, capable of processing diverse imaging modalities while providing spatially reliable prediction estimates for clinical practice. Nevertheless, existing vision-language model solutions suffer from poor domain generalization and exhibit limited capability for uncertainty estimation to model the cumulative unreliability of cross-modal alignment across network layers. In this paper, we propose UniMedSeg, a unified vision-language framework for multi-modal medical image segmentation built upon frozen pre-trained vision-language backbones. Our framework incorporates modality-aware adaptation, cross-layer uncertainty-guided encoding, and a frequency-spatial dual-prompt decoder, wherein the progressive uncertainty propagation mechanism of the encoder suppresses misleading cross-modal features, whereas the decoder disentangles anatomical structure and boundary details in the frequency domain under dual textual prompts. Auxiliary losses further sharpen boundary representations and calibrate pixel-wise uncertainty maps. Extensive experiments demonstrate that UniMedSeg preserves competitive in-domain segmentation performance, improves domain generalization, and yields clinically useful spatial risk estimation. The source code is available at https://github.com/699zjl/UniMedSeg.

**Highlights:**

- We propose **UniMedSeg**, a unified vision-language model for five-modality medical segmentation that outputs segmentation masks and clinical-oriented pixel-level uncertainty maps.
- A cross-layer uncertainty propagation mechanism is designed to dynamically suppress unreliable vision-language alignment by recursively accumulating historical reliability signals.
- A frequency-spatial dual-prompt decoder disentangles anatomical structure and boundary details via FFT-based separation and branch-specific text-guided attention to reduce frequency-domain artifacts.
- Auxiliary boundary contrastive and uncertainty calibration objectives enhance boundary discrimination and enforce consistency between predicted uncertainty and segmentation errors.
- Extensive experiments on multi-modal medical datasets verify that UniMedSeg yields competitive in-domain performance and favorable out-of-distribution generalization capability.

## 1. Introduction

Medical image segmentation serves as a fundamental task in surgical planning, disease assessment, and long-term follow-up. In recent years, deep learning has significantly improved the performance of organ and lesion segmentation, yet its practical deployment continues to face multifaceted challenges. First, pixel-level annotation typically requires substantial time investment from domain experts and is susceptible to inter-observer variability (Chen et al., 2025; Ma et al., 2024; Schmidt et al., 2023; Liu et al., 2022). Second, boundaries between lesions and normal tissues are often ambiguous with gradual intensity transitions, introducing inherent visual ambiguity; existing deterministic models tend to produce erroneous yet highly confident predictions in such scenarios, lacking clinical risk alerts (Yang et al., 2025b; Wang et al., 2025; Han et al., 2025). Third, models trained on single datasets struggle to generalize to unseen data sources and cannot warn clinicians of out-of-distribution cases (Koleilat et al., 2026; Safdari et al., 2026; Yang et al., 2025a). Under such practical constraints, spatially-localized risk alerting for model predictions bears critical clinical value. Highlighting low-confidence regions guides clinicians toward targeted manual review, mitigating misdiagnosis and missed-diagnosis risks arising from over-reliance on model outputs. This necessitates a unified segmentation framework supporting multi-modal inputs, uncertainty-aware spatial risk estimation, and strong generalization across clinical scenarios.

Deep learning approaches built upon U-Net (Ronneberger et al., 2015) and Transformer architectures (Dosovitskiy et al., 2020) have driven substantial progress in medical image segmentation. However, studies (Mehrtash et al., 2020; Ye et al., 2023; Ao et al., 2023) have shown that such deterministic networks can produce overconfident predictions under challenging conditions, including out-of-distribution (OOD) inputs (Karimi and Gholipour, 2022; Shen et al., 2026), ambiguous tissue boundaries (Cheng et al., 2026; Adams and Elhabian, 2023), and imaging noise. Lacking built-in uncertainty warning mechanisms (Koleilat et al., 2026; Adams and Elhabian, 2023; Rathore et al., 2025), these models propagate unreliable evidence through successive layers without providing spatially localized risk estimates for clinical review. Vision-language models pave the way toward such a unified multi-modal framework, supporting text-guided segmentation across diverse medical imaging modalities (Zhang et al., 2023; Khattak et al., 2024). Pre-training frameworks represented by CLIP (Radford et al., 2021) establish cross-modal semantic alignment through large-scale image-text contrastive learning and leverage text prompts to guide the localization and segmentation of anatomical targets (Lüddecke and Ecker, 2022; Cho et al., 2024). However, complex conditional dependencies between visual evidence and linguistic concepts in medical images (Elallaf et al., 2026), along with cross-modal semantic gaps (Lin et al., 2026; Zhang et al., 2025), limit the direct transferability of pre-trained models. Existing adaptation methods employ parameter-efficient strategies including prompt learning (Zhou et al., 2022; Koleilat et al., 2025a), low-rank adaptation (Zanella and Ben Ayed, 2024), and lightweight adapters (Dhakal et al., 2024; Gao et al., 2024), effectively improving cross-domain generalization (Pan et al., 2025). Nevertheless, these approaches primarily focus on enhancing semantic alignment without explicitly modeling how the reliability of cross-modal interactions evolves across spatial locations and encoder layers.

Two intertwined challenges remain unaddressed. First, alignment unreliability accumulates incrementally across encoder layers. Unreliable cross-modal correspondences propagate at each fusion step and degrade downstream representations, yet few existing approaches establish dedicated mechanisms to track or mitigate such cumulative errors. Recent work has enhanced feature reliability through causal intervention (Chen et al., 2024; Yang et al., 2026; Liang et al., 2026), probabilistic modeling (Jha et al., 2024; Upadhyay et al., 2023; Koleilat et al., 2026), and frequency-domain analysis (Yu et al., 2025; Gong et al., 2025). Nevertheless, most of these methods compute reliability locally within individual modules and do not support continuous uncertainty propagation across encoder layers. Consequently, unreliable features cannot be persistently suppressed, and accumulated uncertainty fails to flow into the decoder to guide feature selection or generate calibrated spatial risk maps.

Second, even well aligned cross modal features entangle structural and boundary information that demands differentiated decoding strategies. Such mixed information complicates direct decoding; frequency domain transforms enable explicit disentanglement of these two components Zhang et al. (2026); Ren et al. (2026), offering a promising paradigm to address segmentation challenges caused by diverse lesion morphologies and indistinct boundaries Song et al. (2026).

Nevertheless, such decomposition works merely at the signal level: low frequency components are contaminated by task irrelevant slow varying artifacts (e.g., luminance offsets), whereas high frequency components suffer from noise, off target textures and spurious edges. Without task driven cues, it cannot suppress intra branch spurious responses, leading to sustained misalignment between decomposed features and segmentation objectives. To alleviate this drawback, existing methods Lüddecke and Ecker (2022); Yu et al. (2025) integrate textual information. However, these textual signals cannot filter or modulate frequency components, making differentiated suppression of low frequency background interference and high frequency noisy textures unattainable. This limitation prevents coordinated improvements in structural integrity and boundary precision.

To address these problems, we present UniMedSeg, a unified framework built upon frozen UniMedCLIP and BiomedBERT encoders. In summary, the main contributions of this work are summarized as follows:

1. We present **UniMedSeg**, a unified vision-language framework for multi-modal medical image segmentation. Integrating modality-specific adaptation, cross-layer uncertainty-guided encoding and frequency-spatial dual-prompt decoding, our single model handles five imaging modalities and outputs segmentation masks together with clinically calibrated pixel-level uncertainty maps.
2. We propose an uncertainty-guided encoder. It forms a closed-loop cross-layer uncertainty propagation pipeline with uncertainty-guided modulation, PVL adapter, uncertainty extractor and uncertainty memory bank, which suppresses unreliable cross-modal tokens and accumulates layer-wise uncertainty for feature modulation.
3. We design a frequency-spatial dual-prompt decoder. It leverages uncertainty-injected feature extraction, Bayesian precision-weighted cross-layer fusion, FFT-based frequency-spatial decomposition and dual-prompt text-guided cross-attention to disentangle structural and detail representations and suppress spurious frequency-domain responses. Complementarily, boundary contrastive loss and uncertainty calibration loss are utilized to strengthen boundary discrimination and align predicted uncertainty with real segmentation errors.

## 2. Related Work

### 2.1. Vision-Language Models for Medical Image Segmentation

Vision-language models such as BiomedCLIP Zhang et al. (2023), PubMedCLIP Eslami et al. (2021), and UniMedCLIP Khattak et al. (2024) have endowed medical image segmentation with strong semantic alignment through contrastive pre-training on clinical image-text corpora, yet transferring these coarse-grained correspondences to fine-grained pixel-level predictions requires task-specific adaptation. Early attempts such as CLIPSeg Lüddecke and Ecker (2022), DenseCLIP Rao et al. (2022), and ZegCLIP Zhou et al. (2023) map vision-language features into pixel space but incur substantial parameter overhead due to task-specific decoders or region-level contrastive mechanisms. To alleviate this burden, parameter-efficient fine-tuning (PEFT) techniques—prompt learning (e.g., MaPLe Khattak et al. (2023)), adapters (e.g., CLIP-Adapter Gao et al. (2024)), and low-rank updates (e.g., CLIP-LoRA Zanella and Ben Ayed (2024))—have been adopted in medical segmentation, substantially reducing training costs while preserving pretrained spatial modeling capabilities. Complementarily, SAM-derived architectures such as MedSAM Ma et al. (2024) and MedCLIP-SAMv2 Koleilat et al. (2025b) have emerged to enhance cross-domain generalization by mitigating the unreliable semantic grounding of the original SAM in the medical domain. However, under this accuracy-centric paradigm, these methods have paid limited attention to prediction calibration and uncertainty quantification for cross-modal alignment, and fail to capture its cumulative unreliability across network layers. This may hinder models from distinguishing well-aligned from ambiguous regions—a limitation that warrants careful consideration in clinical deployment.

### 2.2 Uncertainty for Vision-Language Models

Deterministic point-embedding in vision-language models (VLMs) cannot model intrinsic input ambiguity and latent distribution shifts, yielding overconfident erroneous predictions on out-of-distribution (OOD) samples. Probabilistic uncertainty estimation provides a principled remedy for improved predictive reliability. Existing probabilistic vision-language modeling paradigms fall into four categories. First, fully trained probabilistic VLMs (e.g., PCME++ Chun (2024)) require full retraining on large-scale datasets and suffer from limited transferability of pre-trained weights. Second, adapter-based methods, including ProbVLM Upadhyay et al. (2023), AsymVLM Ju et al. (2026) and MedCLIPSeg Koleilat et al. (2026), freeze the backbone and only train lightweight adapters to map final encoder features into probabilistic distributions, yet they fail to exploit multi-layer representations, support cross-layer uncertainty propagation, or model cumulative uncertainty during iterative inference. Third, training-free bayesian posterior approaches (e.g., BayesVLM Baumann et al. (2026)) merely estimate the posterior of the final projection layer and discard intermediate-layer features. Fourth, domain-tuned fine-tuning approaches, exemplified by MedProbCLIP Elallaf et al. (2026), require full-encoder fine-tuning using domain-specific paired data, which results in degraded out-of-domain generalizability. To address these limitations, this work proposes a cross-layer uncertainty propagation module to implement cumulative historical uncertainty and bidirectional uncertainty propagation across encoder and decoder layers.

### 2.3. Frequency-Domain Modeling in Visual Representation

Frequency-domain modeling techniques are divided into two categories based on text involvement: pure-visual frequency-domain modeling and multimodal frequency-domain modeling with text input. (1) Pure-visual frequency-domain modeling comprises frequency-decomposition approaches built upon DWT (e.g., WaveFormer Al Hasan et al. (2025), WFDENet Li et al. (2026)), DCT (SpectralMamba-UNet Zhang et al. (2026)), and FFT (PFESA Li et al. (2025)), alongside parameter-efficient fine-tuning methods that embed frequency-aware adapters into pre-trained backbones (e.g., FreqFiT Ly and Nguyen (2025)). These methods rely solely on visual frequency information and exclude textual semantic guidance, making them incapable of incorporating task-aware prior knowledge to adaptively optimize frequency-domain representations. (2) Multimodal frequency-domain modeling methods incorporate textual semantic information for feature optimization. Specifically, CLIPSeg Lüddecke and Ecker (2022) realizes text-driven segmentation with single textual guidance, FMISeg Yu et al. (2025) conducts overall interaction between unified text features and high- and low-frequency branches, while FCPrompt Liu et al. (2025) and FE-CLIP Gong et al. (2025) design prompt paradigms from global/local and normal/abnormal dimensions, respectively. Nevertheless, these methods cannot perform differentiated semantic modulation for low-frequency structural branches and high-frequency detail branches, thus failing to suppress irrelevant interference in the two frequency bands separately.

## 3. Method

### 3.1. Overview

Given a medical image *I* ∈ℝ^*B×C×H×W*^, a text prompt *T* describing the target structure, and a modality identifier *m* ∈{0, 1, 2, 3, 4} corresponding to {ultrasound, mri, x-ray, endoscopy, dermoscopy}, UniMedSeg produces a segmentation mask *ŷ* ∈ℝ^*H×W*^ and an uncertainty map *U* ∈ ℝ^*H×W*^. Fig. 1 depicts this unified vision-language encoder-decoder for multi-modal medical image segmentation. Input images and modality-augmented text prompts are fed into dual encoders with progressive uncertainty propagation (Sec. 3.2), where an uncertainty memory bank accumulates token-wise confidence via uncertainty-guided modulation and PVL-Adapter fusion for progressively refining feature reliability across ambiguous, artifact-prone and shifted input conditions. Across-layer features are passed to the frequency-spatial dual-prompt decoder, whose core components—bayesian precision-weighted aggregation and dual prompt text-guided cross attention—are detailed in Sec. 3.3 to filter frequency-domain information via semantic conditioning and suppress irrelevant interference for precise segmentation

**Figure 1:**
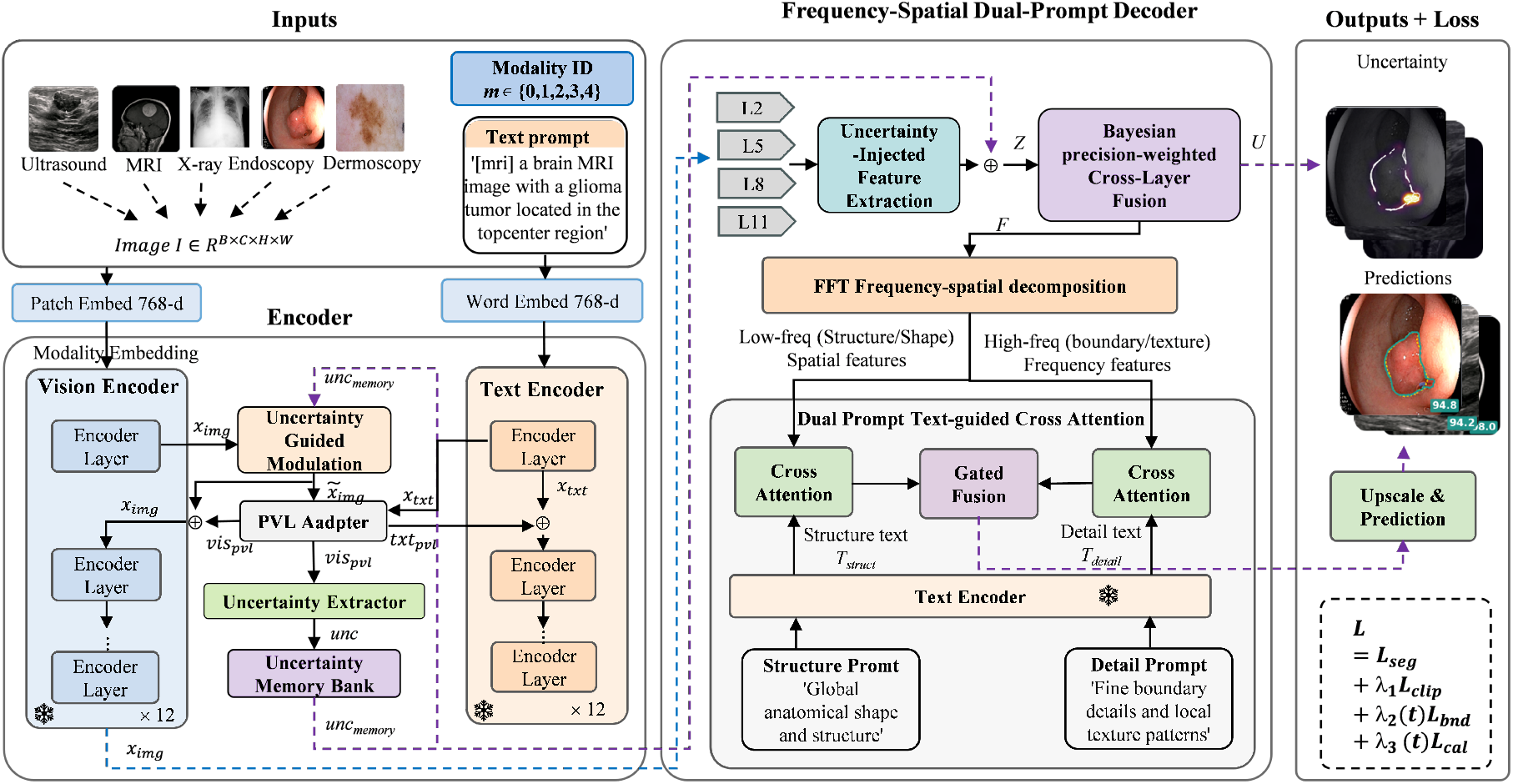
Overview of the proposed UniMedSeg framework. The encoder (left) employs dual vision-text branches with modality embedding injection and progressive uncertainty propagation modules (Uncertainty Guided Modulation, PVL Adapter, Uncertainty Extractor, and Uncertainty Memory Bank). The decoder (right) aggregates uncertainty-injected feature representations with bayesian precision-weighted cross-layer fusion, then decomposes the fused representation via FFT into spatial and frequency branches, each guided by specialized text prompts through cross-attention. Frozen modules are marked with ❄.

**Figure 2:**
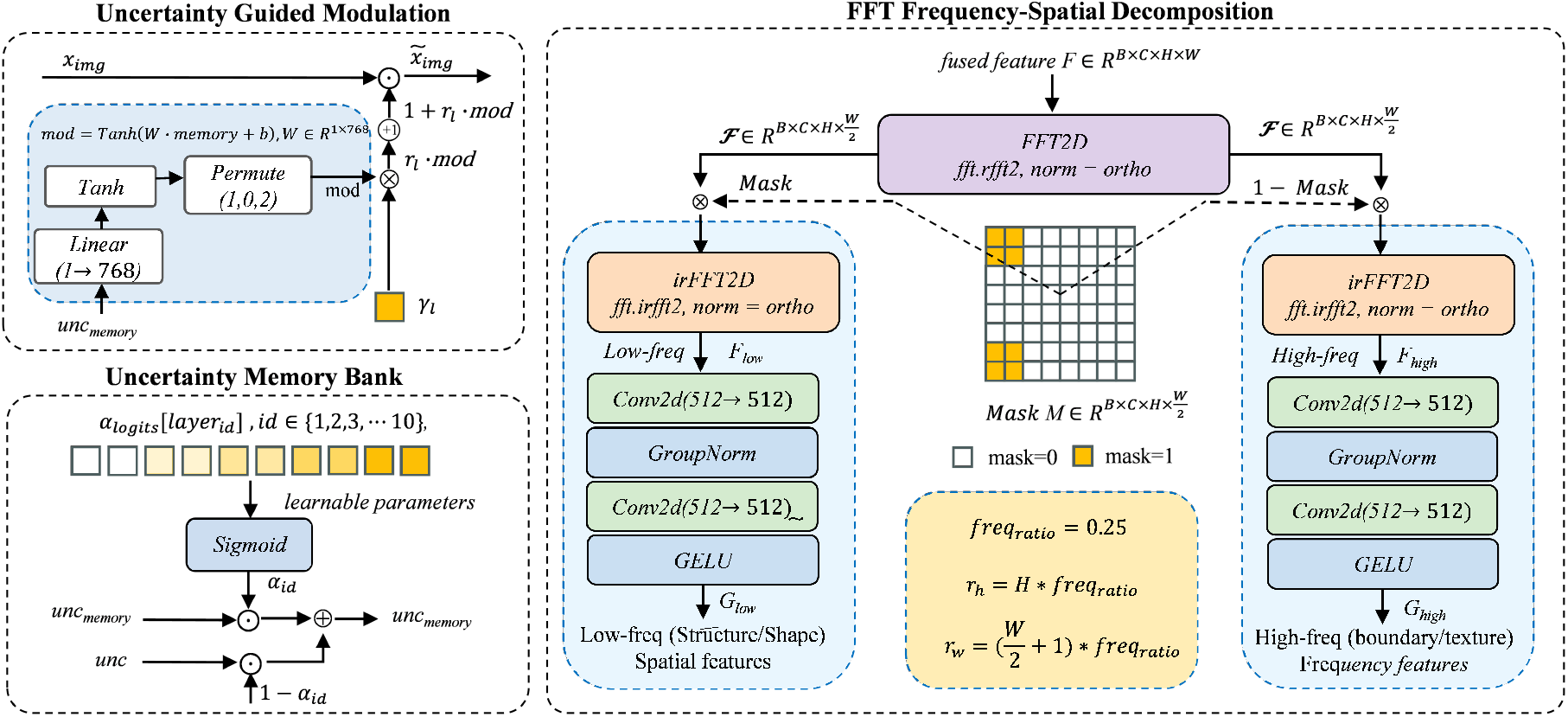
Detailed illustration of key modules. (a) Uncertainty Guided Modulation: accumulated uncertainty memory modulates visual tokens through a linear projection (1 → 768), Tanh activation, and learnable scaling *γ*_*l*_ before each PVL adapter. (b) Uncertainty Memory Bank: per-layer learnable weights *α*_*l*_ control exponential moving average accumulation of uncertainty signals across 10 fusion layers. (c) FFT Frequency-Spatial Decomposition: fused features are transformed to the frequency domain via 2D real FFT, then separated into low-frequency (structure/shape) and high-frequency (boundary/texture) components by a binary mask (*r* = 0.25), each processed by depthwise separable convolutions.

### 3.2. Uncertainty-Guided Encoder

#### 3.2.1. Vision-Language Backbone

We employ the frozen UniMedCLIP (ViT-B/16) Khattak et al. (2024) vision encoder and BiomedBERT Zhang et al. (2023) text encoder as dual backbones, producing visual token sequences *x*_img_ ∈ℝ^*B×*(*N* +1)*×d*^ (*N* = 196, *d* = 768) and text embeddings *x*_txt_ ∈ℝ^*B×L×d*^. To make the model aware of modality-specific visual statistics, each anatomical text prompt is prepended with a modality tag (e.g., “[ultrasound] breast lesion”). A learnable modality embedding indexed by *m*∈ {0, 1, 2, 3, 4} is injected into the visual token sequence:

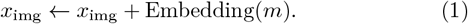

The frozen encoders preserve pre-learned cross-modal alignment, whereas the trainable modality embeddings enable adaptive feature modulation across heterogeneous imaging modalities.

#### 3.2.2. Progressive Uncertainty Propagation

To continuously suppress unreliable cross-modal alignment features in encoding, we model uncertainty as layer-propagated state memory. At the *l*-th fusion stage, the model proceeds through a four-step cycle. It first modulates current visual features with previously accumulated uncertainty memory, then conducts vision-language interaction via PVL-Adapter, followed by extracting new token-wise uncertainty from visual residuals, and finally aggregates the derived uncertainty into memory for subsequent layers. This establishes a strict causal propagation chain: historical uncertainty feedback → visual modulation → PVL fusion → uncertainty extraction → memory update. Driven by this cross-layer feedback loop, the model progressively refines feature reliability throughout encoding and yields spatially localized risk alerts for clinical deployment.

1. **Uncertainty Guided Modulation**. Let 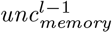 denote the per-token uncertainty memory accumulated before the *l*-th stage, with shape *B ×* (*N* + 1) *×* 1. It is mapped to the visual feature dimension via a learnable linear layer with Tanh:

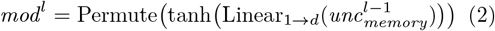

where *d* = 768. Before the current visual features enter the PVL Adapter, element-wise multiplicative modulation is applied:

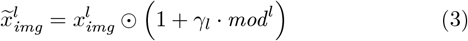

where ⊙ denotes element-wise multiplication and *γ*_*l*_ is a learnable per-layer scaling factor initialized to 0.01. The historical uncertainty memory encodes the reliability of cross-modal alignment, instead of measuring the intrinsic quality of visual features. Concretely, a learnable linear projection produces channel-wise modulation weights for every token, which are used to selectively reweight 768-dimensional visual features. Tokens with poor alignment histories are attenuated, whereas reliably aligned tokens are boosted. This mechanism dynamically calibrates visual feature responses before cross-modal fusion takes place.
2. **PVL Adapter**. The PVL Adapter (Koleilat et al., 2026) fuses the modulated visual features and text features through bidirectional probabilistic cross-attention. It first projects both modalities into a shared low-dimensional space *d*_*a*_ via learnable down-projections, then performs bidirectional cross-modal interaction where visual and text tokens mutually attend to each other via probabilistic attention:

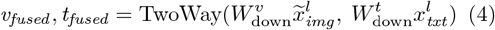

and finally projects the fused representations back to their original dimensions:

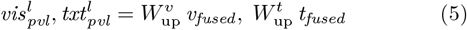 These outputs are added back to the modulated inputs via residual connections:

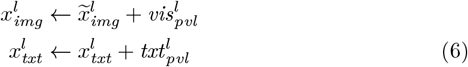 The TwoWay module executes bidirectional cross-modal interaction via two cascaded probabilistic cross-attention layers: the first layer uses visual tokens as Q and text tokens as K/V, while the second layer reverses the roles. Within each layer, the query projection outputs single-channel features, whereas both key and value projections output double-channel features—the first half serves as the mean (*K*_*µ*_, *V*_*µ*_) and the second half, after Softplus activation, as the variance (*K*_var_, *V*_var_)—explicitly modeling cross-modal alignment uncertainty as probabilistic distributions. Given these probabilistic representations, attention scores incorporate a variance penalty beyond semantic similarity:

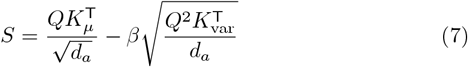

where *Q* is the query, *K*_*µ*_ and *K*_var_ are the mean and variance of keys. The first term measures cross-modal semantic affinity; the second down-weights high-variance keys, suppressing unreliable context without manual thresholding. After computing attention weights *α* = softmax(*S*), Values are sampled from their probabilistic distributions and aggregated:

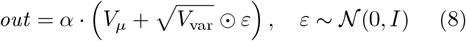

where *V*_*µ*_ and *V*_var_ are the mean and variance of values. The aggregated output is projected and fused with the original query via a learnable gate *g* ∈ (0, 1):

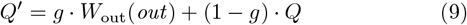

where *Q*^*′*^ denotes the updated query, which corresponds to *v*_*fused*_ and *t*_*fused*_ in Eq. (4) after bidirectional execution. The two modules operate sequentially within each fusion stage. Historical uncertainty first adjusts the visual representation before cross-modal fusion, while probabilistic attention evaluates the reliability of the current visual-text interaction during fusion. The resulting visual residual is then passed to the uncertainty extractor to produce the uncertainty state for subsequent stages.
3. **Uncertainty Extractor**. The visual residual 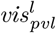 produced by the PVL adapter represents the outcome of the current visual-text interaction. We use this residual as input to a lightweight per-token predictor that estimates the corresponding uncertainty state:

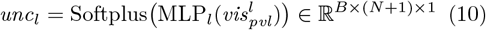

where MLP_*l*_ : ℝ^768^ → ℝ^128^ → ℝ^1^ with GELU activation. The Softplus activation constrains the output to be non-negative, as required for subsequent aggregation and precision-weighted decoding. Here, *unc*_*l*_ denotes a learned token-wise uncertainty score rather than the variance derived from an explicit posterior distribution. It receives no explicit constraint to match ground-truth segmentation error; instead, this property is induced by the uncertainty calibration objective (Sec. 3.4): higher uncertainty scores tend to coincide with larger segmentation errors. Produced by the uncertainty extractor, *unc*_*l*_ yields deterministic token-level outputs without stochastic sampling during inference. This signal is propagated across successive fusion stages via the memory bank to modulate down-stream visual representations.
4. **Uncertainty Memory Bank**. Uncertainty scores from an individual fusion stage capture only layer-local cues and lack holistic cross-stage awareness. To aggregate cross-stage uncertainty and exploit historical reliability statistics for subsequent fusion steps, we define *unc*_*memory*_, a forward-updated uncertainty state that accumulates token-level scores across fusion stages via a learnable exponential moving average. Recursively updated throughout one forward pass, this state is maintained independently for each input sample. At each fusion stage *l*, the memory state is updated as:

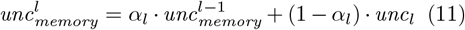

where *α*_*l*_ = *σ*(*w*_*l*_) ∈ (0, 1) denotes a stage-specific learnable weight and *σ*(·) is the sigmoid function. The weight *α*_*l*_ governs the balance between historical accumulated evidence and newly arrived stage-wise estimates: larger *α*_*l*_ preserves information from preceding layers, while smaller values place greater emphasis on current observations. Through this adaptive weighting, the model preserves long-range uncertainty priors while continuously fusing up-to-date cross-modal cues.

### 3.3. Frequency-Spatial Dual-Prompt Decoder

Encoder layers provide complementary semantics but differ in reliability. The encoder produces token-level uncertainty characterizing cross-modal alignment quality within individual layers, whereas effective cross-layer fusion requires spatial uncertainty that quantifies the relative reliability of different layers at each position. We introduce an uncertainty-injected feature extraction module to bridge these two stages: it first incorporates encoder-accumulated uncertainty into multi-level visual representations, then derives layer-specific spatial uncertainty maps for bayesian precision-weighted fusion. Since direct frequency decomposition may retain task-irrelevant noise, we further decouple the fused features into low-frequency structural and high-frequency detail components via FFT, and apply paired structural and detail prompts to regularize both branches. This design preserves feature complementarity while sharpening semantic focus on target regions and ambiguous boundaries.

#### 3.3.1. Uncertainty-Injected Feature Extraction

To establish a complete reliability propagation path from the encoder to the decoder, we explicitly inject the cross-modal alignment information 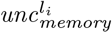 accumulated in the encoder into the visual representation 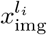. Specifically, two separate projection layers are adopted to perform feature mapping and channel-dimension alignment for the visual representation and uncertainty state respectively, followed by element-wise addition:

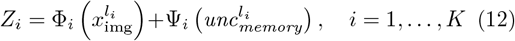

where Φ_*i*_ denotes the visual projection for the *i*-th encoder feature, implemented by a 1 *×* 1 convolution followed by GroupNorm and GELU. Ψ_*i*_ projects the corresponding uncertainty prior into the same channel dimension using a 1 *×* 1 convolution. The projected visual feature and uncertainty prior are then added to form *Z*_*i*_. This operation injects encoder-side cross-modal reliability information into each multi-scale representation, allowing the subsequent uncertainty prediction head to estimate the task-aware spatial reliability of that scale for cross-layer fusion.

#### 3.3.2. Bayesian Precision-Weighted Cross-Layer Fusion

To quantify the spatial reliability of each layer representation *Z*_*i*_, we employ a dedicated uncertainty prediction head. Specifically, for the *i*-th representation *Z*_*i*_, we estimate its spatial uncertainty through an independent uncertainty prediction head:

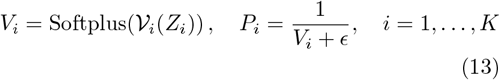

where *V*_*i*_ denotes the uncertainty prediction head corresponding to the *i*-th representation, composed of a 3 *×* 3 convolution, GELU activation, and a 1 *×* 1 convolution. The Softplus ensures the uncertainty map *V*_*i*_ is non-negative, and *ϵ* is a small constant added to the denominator. The precision weight *P*_*i*_ converts uncertainty to precision: higher uncertainty *V*_*i*_ yields lower precision *P*_*i*_, thus reducing the layer’s contribution to fusion, while lower uncertainty yields higher precision, increasing its influence.

Based on the precision weights of each layer, the fused representation *F* and its uncertainty estimate *U* are computed as:

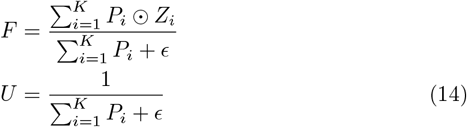

Here, ⊙ denotes element-wise multiplication. The precision weighting mechanism dynamically modulates the contribution of each encoder layer according to local representation reliability: high-precision features dominate the fusion process, while low-precision features are adaptively attenuated. Meanwhile, *U*, defined as the reciprocal of the accumulated precision, characterizes the overall uncertainty of the fused representation. This yields the fused feature *F* and uncertainty *U*, where *F* feeds into frequency-domain decomposition.

#### 3.3.3. FFT Frequency-Spatial Decomposition

The fused representation *F* entangles structural content (low-frequency) and boundary details (high-frequency), making single-pathway processing prone to over-smoothing or boundary artifacts. We decouple these components via FFT and refine them independently:

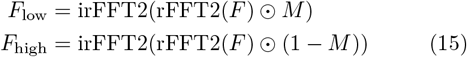

where rFFT2(·) and irFFT2(·) denote the two-dimensional real-valued fast Fourier transform and its inverse, and *M* is a binary mask in the frequency domain. *F*_low_ preserves the overall structural variations of the target region, while *F*_high_ highlights local boundary and texture variations.

Subsequently, the two frequency components are processed through independent convolutional branches:

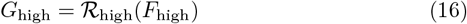

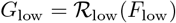

where *R*_low_ and *R*_high_ denote the low-frequency and high-frequency convolutional branches, respectively. Each branch consists of a 3 *×* 3 depthwise convolution, GroupNorm, GELU activation, and a 1 *×* 1 convolution, extracting spatial patterns and reorganizing channel information.

#### 3.3.4. Dual Prompt Text-Guided Cross Attention

To inject task-specific semantic priors into the decomposed frequency branches, we encode paired textual prompts describing anatomical structures and boundary details via BiomedBERT into *T*_struct_ and *T*_detail_, then apply branch-specific cross-attention:

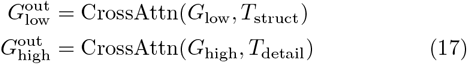

where visual features serve as queries and prompt features serve as keys and values, with residual connections preserving the original frequency characteristics.

The two semantically guided branches are then combined via learnable gating:

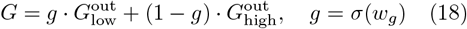

where *w*_*g*_ is a learnable scalar that adaptively balances structural and detail contributions. The fused representation *G* is upsampled and projected to produce segmentation logits, while the decoder uncertainty *U* from Eq. 14 is interpolated to the target resolution as the pixel-level uncertainty map.

### 3.4. Training Strategy

The model is trained with the following composite objective:

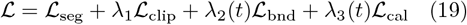

where *L*_seg_ is the segmentation loss, *L*_clip, bnd_, and *L*_cal_ are image-text alignment, boundary contrastive, and uncertainty constraint losses, respectively. The weights *λ*_2_(*t*) and *λ*_3_(*t*) are gradually increased during training for stable optimization. Detailed hyperparameter settings are provided in Section 4.1.

1. **Segmentation loss**. To balance region-level overlap and pixel-level classification, the segmentation loss combines Dice loss and Binary Cross-Entropy:

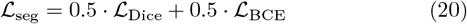
2. **Image-text alignment loss**. Let 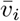 and 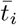 denote the normalized visual and text representations of the *i*-th sample, where the visual representation is aggregated from patch features. We adopt a bidirectional soft contrastive objective:

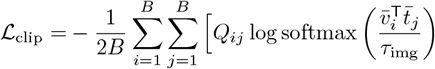

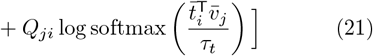

where *B* is the batch size, 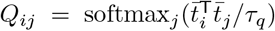 is the soft target distribution derived from pairwise text similarities, with *τ*_*q*_ controlling the sharpness of the soft labels, and *τ*_img_, *τ*_*t*_ are temperature parameters for image-to-text and text-to-image prediction distributions. This loss constrains image-text matching in both directions while preserving flexibility between semantically similar texts.
3. **Boundary contrastive loss**. We map the ground-truth mask to a 14 *×* 14 patch grid and apply morphological operations to obtain boundary-foreground patch set *A* (anchors), interior-foreground patches, and boundary-background patch set *N* (negatives). The positive set *P* includes interior-foreground patches and the corresponding text representation:

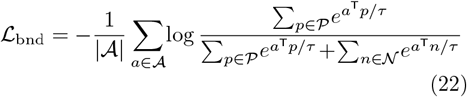

where *τ* is the boundary contrastive temperature, |*A*| is the number of anchor patches, *P* includes interior-foreground patches and text representation, and *N* comprises boundary-background patches. This loss encourages boundary-foreground representations to align with interior-foreground and text semantics while staying away from boundary-background, thereby enhancing discriminability in boundary regions.
4. **Uncertainty calibration loss**. To align decoder-predicted uncertainty with actual prediction errors, we define:

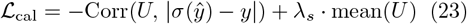

where *U* is the fused uncertainty from Eq. 14 interpolated to the target resolution, *ŷ* denotes the segmentation logits, *y* is the ground-truth mask, and *σ*(·) is the sigmoid function. The correlation term encourages high-uncertainty regions to correspond to large prediction errors, while the mean constraint (with weight *λ*_*s*_ = 0.01) limits the overall magnitude of uncertainty to prevent indiscriminately high outputs.

## 4. Experiments

### 4.1. Experimental Setup

#### 4.1.1. Datasets

Following Koleilat et al. (2026), we conduct evaluations over 16 medical image segmentation datasets covering 5 imaging modalities (ultrasound, MRI, dermoscopy, endoscopy, X-ray) and 6 organs (breast, brain, skin, colon, chest, pancreas). The benchmark contains 6 source datasets (BUSI, BTMRI, ISIC, Kvasir-SEG, QaTa-COV19, EUS) for training and in-domain evaluation, and 10 out-of-distribution (OOD) datasets (BUSUC, BUSBRA, BUID, UDIAT, CVC-ColonDB, CVC-ClinicDB, CVC-300, BKAI, BRISC, UWaterlooSkinCancer) for domain-generalization tests.

#### 4.1.2. Evaluation Protocol

Three groups of experiments are designed: (1) **Data-efficiency evaluation**: We train models with 10%, 25%, 50%, and 100% fractions of training samples from the 6 source datasets, and report the averaged DSC and NSD metrics computed over these six datasets. (2) **Domain generalization**: Domain-specific models are trained on one source dataset and directly tested on corresponding unseen OOD target datasets without any adaptation. (3) **Unified segmentation**: A single model is jointly trained on 15 datasets with EUS held out as an unseen target (15DS), or on all 16 datasets (16DS), and compared with domain-specific models trained on four clinical cohorts: Breast Ultrasound (5 datasets), Polyp Endoscopy (5 datasets), Brain MRI (2 datasets), and Skin Dermoscopy (2 datasets).

#### 4.1.3. Implementation Details

All models are trained for 100 epochs using AdamW optimizer (lr=2 *×* 10^−4^, weight decay=0.01) with cosine annealing (*η*_min_ = 10^−5^), batch size 16, and gradient clipping at 1.0. The segmentation loss combines Dice and BCE with equal weights (0.5 each). For the unified 16DS model, loss weights are *λ*_1_ = 0.1, *λ*_2_(*t*) = 0.3 · min(*t/*5, 1), and *λ*_3_(*t*) = 0.05 · min(*t/*10, 1). For domain-specific models, *λ*_2_(*t*) = 0.15 · min(*t/*20, 1) and *λ*_3_(*t*) = 0.05 · min(*t/*30, 1) with longer warmup. Temperature parameters are *τ*_img_ = *τ*_*t*_ = *τ*_*Q*_ = 0.07 and *τ* = 0.07 for boundary contrastive learning. Experiments run on a single NVIDIA RTX 5090 GPU.

### 4.2. Data Efficiency Evaluation

As shown in Table 1, UniMedSeg demonstrates superior data efficiency across all training regimes, consistently surpassing both traditional unimodal architectures and recent vision-language baselines. At the most challenging 10% data regime, UniMedSeg achieves 85.25% DSC, outperforming the competitive CLIP-based MedCLIPSeg by **+4.15** percentage points (pp) and CAT-Seg by **+6.49 pp**. This advantage persists at 25% data with **+1.60 pp** and **+5.56 pp** gains over MedCLIPSeg and CAT-Seg, respectively. These improvements validate that our progressive uncertainty propagation and frequency-spatial dual-prompt decoder enable more effective feature refinement and uncertainty-aware learning from limited annotations.

**Table 1.** Data-efficiency evaluation. Average DSC and NSD (%) for segmentation methods with varied training-data fractions. Bold = best, underlined = second-best performance.

| Method | 10% Data |  | 25% Data |  | 50% Data |  | 100% Data |  |
| --- | --- | --- | --- | --- | --- | --- | --- | --- |
|  | DSC ↑ | NSD ↑ | DSC ↑ | NSD ↑ | DSC ↑ | NSD ↑ | DSC ↑ | NSD ↑ |
| Unimodal Approaches |  |  |  |  |  |  |  |  |
| UNet Ronneberger et al. (2015) | 60.95 | 64.43 | 62.74 | 66.16 | 71.61 | 75.14 | 78.49 | 82.07 |
| UNet++ Zhou et al. (2019) | 63.72 | 67.08 | 65.86 | 69.21 | 73.15 | 76.31 | 78.44 | 81.79 |
| AttnUNet Oktay et al. (2018) | 62.78 | 66.25 | 64.97 | 68.53 | 71.34 | 74.96 | 76.30 | 79.77 |
| nnUNet Isensee et al. (2021) | 73.45 | 77.37 | 76.73 | 80.66 | 78.86 | 82.68 | 81.40 | 85.08 |
| Swin-UNet Cao et al. (2022) | 53.04 | 57.91 | 54.69 | 59.24 | 55.89 | 61.25 | 65.03 | 69.32 |
| TransUNet Chen et al. (2021) | 52.69 | 56.38 | 55.25 | 58.95 | 55.22 | 59.30 | 67.22 | 71.15 |
| Generic Text-driven Approaches |  |  |  |  |  |  |  |  |
| LViT Li et al. (2023) | 66.51 | 68.80 | 75.66 | 78.12 | 78.88 | 81.34 | 83.35 | 85.89 |
| Ariadne's ThreadZhong et al. (2023) | 61.34 | 62.75 | 63.09 | 64.51 | 65.65 | 66.92 | 70.07 | 71.49 |
| CLIP-Based Approaches |  |  |  |  |  |  |  |  |
| CLIPSeg Lüddecke and Ecker (2022) | 74.66 | 77.75 | 78.31 | 81.34 | 79.63 | 82.58 | 84.87 | 87.74 |
| DenseCLIP Rao et al. (2022) | 67.84 | 70.33 | 70.23 | 72.70 | 72.09 | 74.45 | 74.19 | 76.89 |
| ZegCLIP Zhou et al. (2023) | 61.25 | 63.72 | 72.46 | 75.01 | 76.21 | 78.80 | 78.98 | 81.69 |
| SAN Xu et al. (2023) | 74.13 | 76.97 | 76.13 | 78.91 | 78.80 | 81.52 | 81.62 | 84.35 |
| MaPLe Khattak et al. (2023) | 66.27 | 68.75 | 71.53 | 73.95 | 74.60 | 77.12 | 74.60 | 77.10 |
| MaPLe + Decoder Khattak et al. (2023) | 74.81 | 77.90 | 79.64 | 82.60 | 82.81 | 85.80 | 84.94 | 87.91 |
| VLSM-Adapter Dhakal et al. (2024) | 74.47 | 77.50 | 77.63 | 80.53 | 80.83 | 83.77 | 83.85 | 86.72 |
| CausalCLIPSeg Chen et al. (2024) | 71.19 | 73.74 | 75.42 | 78.00 | 78.60 | 81.22 | 81.34 | 84.20 |
| CAT-Seg Cho et al. (2024) | 78.76 | 81.50 | 81.12 | 83.92 | 83.32 | 85.61 | 85.90 | 88.31 |
| MedCLIPSeg Koleilat et al. (2026) | <u>81.10</u> | <u>83.94</u> | <u>85.08</u> | <u>87.85</u> | <u>87.18</u> | <u>89.95</u> | <u>88.66</u> | <u>91.35</u> |
| **UniMedSeg (Ours)** | **85.25** | **88.25** | **86.68** | **89.74** | **87.67** | **90.70** | **88.68** | **91.68** |

### 4.3. Domain Generalization

We evaluate domain generalization on breast-ultrasound, polyp endoscopy, brain MRI, and skin dermoscopy tasks. The domain-specific UniMedSeg model is trained independently for each clinical domain, whereas UniMedSeg (6DS) is jointly trained on BUSI, BTMRI, ISIC, Kvasir, QaTa-COV19, and EUS. Results are presented in Tables 2 and 3. The domain-specific UniMedSeg yields an average in-distribution (ID) DSC of 89.55%, compared with 89.35% for UniMedSeg (6DS). The marginal gap of 0.21 pp demonstrates that joint training achieves source-domain performance comparable to domain-specific training. This is attributed to modality embedding, which preserves modality-specific features while learning cross-dataset shared representations.

**Table 2.** Domain generalization (DSC %). Part 1: **bold**/<u>underlined</u>: best and runner-up results for domain-specific models trained on a single source dataset, respectively. Purple entries indicate universal segmentation models outperforming all domain-specific models trained on all source datasets (BUSI, BTMRI, ISIC, Kvasir, QaTa-COV19, EUS). All entries are single runs with seed 666.

| Method | Breast Ultrasound |  |  |  |  | Polyp Endoscopy |  |  |  |  |
| --- | --- | --- | --- | --- | --- | --- | --- | --- | --- | --- |
|  | Source | Target |  |  |  | Source | Target |  |  |  |
|  | BUSI | BUSBRA | BUSUC | BUID | UDIAT | Kvasir | ColonDB | ClinicDB | CVC300 | BKAI |
| *Domain-specific training:* |  |  |  |  |  |  |  |  |  |  |
| LViT<sup>†</sup> Li et al. (2023) | 75.32 | 59.41 | 67.95 | 53.51 | 65.60 | 85.29 | 60.01 | 75.27 | 70.22 | 67.17 |
| CLIPSeg<sup>†</sup> Lüddecke and Ecker (2022) | 80.95 | 63.66 | 75.03 | 68.43 | 56.67 | 81.98 | 59.93 | 71.49 | 72.74 | 66.46 |
| DenseCLIP<sup>†</sup> Rao et al. (2022) | 71.85 | 53.34 | 70.97 | 63.53 | 54.93 | 79.32 | 56.38 | 68.08 | 64.71 | 61.63 |
| ZegCLIP<sup>†</sup> Zhou et al. (2023) | 72.08 | 61.08 | 73.57 | 71.75 | 52.41 | 78.46 | 53.46 | 69.75 | 60.73 | 65.60 |
| SAN<sup>†</sup> Xu et al. (2023) | 77.99 | 64.37 | 74.15 | 58.13 | 61.98 | 83.16 | 61.82 | 74.46 | 80.36 | 69.31 |
| MaPLE<sup>†</sup> Khattak et al. (2023) | 66.37 | 50.08 | 71.52 | 70.77 | 57.81 | 76.12 | 48.09 | 59.64 | 63.80 | 56.94 |
| MaPLE+Dec<sup>†</sup> Khattak et al. (2023) | 80.49 | 55.89 | 64.96 | 60.66 | 59.44 | 83.46 | 61.53 | 71.20 | 74.62 | 66.93 |
| VLSM-Adp<sup>†</sup> Dhakal et al. (2024) | 80.90 | 68.48 | 82.37 | 75.26 | 69.16 | 85.89 | 63.51 | 76.09 | 75.24 | 71.59 |
| CausalCLIP<sup>†</sup> Chen et al. (2024) | 76.11 | 55.87 | 69.12 | 64.49 | 48.90 | 78.77 | 41.65 | 57.54 | 45.77 | 52.56 |
| CAT-Seg<sup>†</sup> Cho et al. (2024) | 81.83 | 70.94 | 81.48 | 73.37 | 70.30 | 86.43 | 68.49 | 70.35 | 78.12 | 74.35 |
| MedCLIPSeg<sup>†</sup> Koleilat et al. (2026) | <u>85.72</u> | **75.06** | <u>84.37</u> | <u>78.99</u> | <u>74.64</u> | **90.15** | <u>71.90</u> | <u>80.80</u> | <u>80.82</u> | <u>79.15</u> |
| **UniMedSeg** | **85.92** | <u>74.46</u> | **85.16** | **80.48** | **78.31** | <u>90.11</u> | **75.46** | **84.21** | **82.32** | **81.46** |
| *Unified training (all source datasets):* |  |  |  |  |  |  |  |  |  |  |
| UniMedSeg (6DS) | 85.66 | 77.47 | 86.89 | 82.53 | 79.10 | 90.26 | 73.43 | 84.13 | 84.42 | 76.69 |

**Table 3.** Domain generalization (DSC %). Part 2: **bold**/<u>underlined</u>: best and runner-up results for domain-specific models trained on a single source dataset, respectively. Purple entries indicate universal segmentation models outperforming all domain-specific models trained on all source datasets (BUSI, BTMRI, ISIC, Kvasir, QaTa-COV19, EUS). All entries are single runs with seed 666.

| Method | Brain MRI |  | Skin Dermatology |  |
| --- | --- | --- | --- | --- |
|  | Source | Target | Source | Target |
|  | BTMRI | BRISC | ISIC | UWaterloo |
| *Domain-specific training:* |  |  |  |  |
| LViT<sup>†</sup> Li et al. (2023) | 81.41 | 71.86 | 91.21 | 58.87 |
| CLIPSeg<sup>†</sup> Lüddecke and Ecker (2022) | 86.33 | 77.61 | 90.55 | 80.19 |
| DenseCLIP<sup>†</sup> Rao et al. (2022) | 70.30 | 34.12 | 89.29 | 53.39 |
| ZegCLIP<sup>†</sup> Zhou et al. (2023) | 76.65 | 66.31 | 81.45 | 38.60 |
| SAN<sup>†</sup> Xu et al. (2023) | 85.27 | 71.60 | 91.39 | 82.51 |
| MaPLE<sup>†</sup> Khattak et al. (2023) | 75.40 | 45.19 | 88.31 | 69.12 |
| MaPLE+Dec<sup>†</sup> Khattak et al. (2023) | 85.08 | 71.46 | 90.10 | 81.83 |
| VLSM-Adp<sup>†</sup> Dhakal et al. (2024) | 85.03 | 68.92 | 91.30 | 82.17 |
| CausalCLIP<sup>†</sup> Chen et al. (2024) | 81.71 | 53.96 | 89.47 | 48.73 |
| CAT-Seg<sup>†</sup> Cho et al. (2024) | 84.86 | 76.28 | 91.27 | 82.02 |
| MedCLIPSeg<sup>†</sup> Koleilat et al. (2026) | <u>88.03</u> | <u>80.92</u> | **92.54** | <u>83.53</u> |
| **UniMedSeg** | **89.69** | **82.44** | <u>92.49</u> | **86.28** |
| *Unified training (all source datasets):* |  |  |  |  |
| UniMedSeg (6DS) | 89.57 | 82.67 | 91.89 | 86.92 |

The unified model exhibits more prominent advantages under out-of-distribution (OOD) settings. Its average target-domain DSC reaches 82.69% across four clinical domains versus 82.30% for the domain-specific model. Over all ten target datasets, the respective OOD averages are 81.43% and 81.06%. UniMedSeg (6DS) surpasses the domain-specific counterpart on 7 out of 10 target datasets, yielding gains of 3.01, 2.05, and 2.10 pp on BUSBRA, BUID, and CVC300. These improvements indicate that cross-domain training boosts representation robustness against source-target distribution shifts, where cross-layer uncertainty propagation suppresses domain-shift-triggered spurious features and the frequency-spatial dual-prompt decoder disentangles structural information from local texture and boundary details.

Nevertheless, the unified model yields performance drops of 2.03, 0.08, and 4.77 pp on ColonDB, ClinicDB, and BKAI, implying that domain-specific training is preferable for datasets with unique image statistics tied to specific acquisition distributions. In comparison, shared knowledge delivers OOD improvements of 1.90, 0.23, and 0.64 pp at the domain level for breast ultrasound, brain MRI, and skin dermoscopy, respectively. Boundary contrastive sharpening further boosts these gains by improving feature discriminability in ambiguous transition regions.

### 4.4. Unified Segmentation Capability

We examine whether a single model jointly trained on heterogeneous modalities can retain in-distribution (ID) accuracy while generalizing to unseen data. UniMedSeg and MedCLIPSeg are trained with identical pipelines under three strategies, each with 25%, 50%, and 100% of the training data: (i) *domain-specific specialists*, trained separately in four clinical domains (BreastUS 5DS, PolypEndo 5DS, BrainMRI 2DS, and SkinDerm 2DS), with ID-DSC averaged over source datasets and OOD-DSC evaluated across modalities; (ii) *Unified-15DS*, jointly trained on 15 datasets with EUS held out; and (iii) *Unified-16DS*, jointly trained and evaluated in-distribution on all 16 datasets. ID and OOD performance are summarized by their harmonic mean, HM = 2 · ID · OOD*/*(ID + OOD). Results are shown in Figure 3.

**Figure 3:**
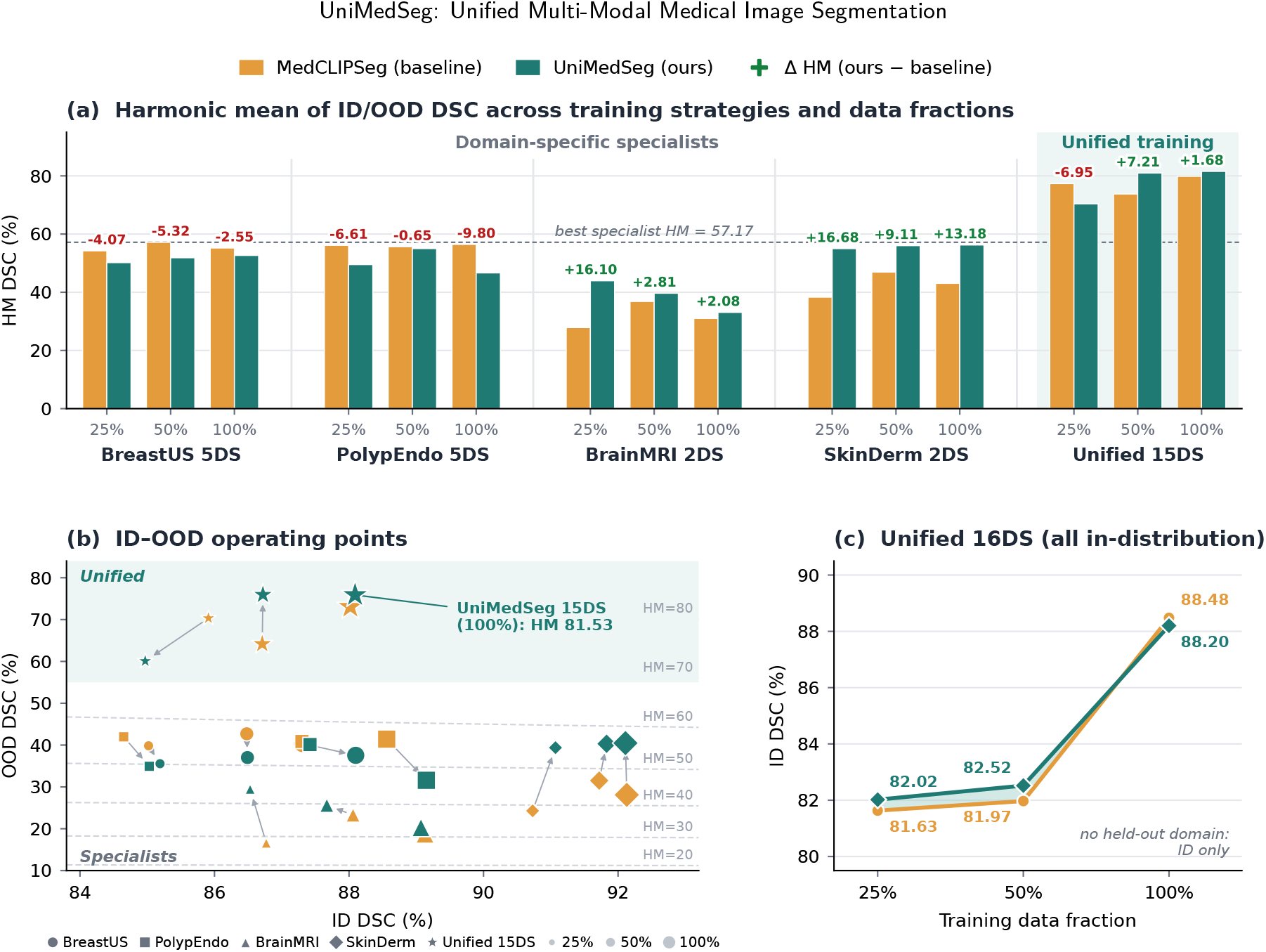
Unified models versus domain-specific models for MedCLIPSeg (orange) and UniMedSeg (teal). (a) HM DSC for each training strategy at 25%, 50%, and 100% training data; numbers above each pair give the HM change of UniMedSeg over MedCLIPSeg, and the dashed line marks the best domain-specific HM (57.17%). (b) ID–OOD operating points; marker shape denotes the training strategy, marker size the data fraction, and arrows point from MedCLIPSeg to UniMedSeg under the same setting. Dashed curves are iso-HM contours. (c) ID DSC of the unified 16DS model, where all domains are in-distribution and OOD/HM are undefined.

#### 4.4.1. Domain-Specific Specialists

Across the four specialist settings, ID-DSC ranges from 84.65% to 92.13% and OOD-DSC from 16.59% to 42.70%, yielding HM values up to 57.17% (dashed line in Figure 3(a)); all specialist operating points fall below the HM = 60 contour in Figure 3(b). Comparing UniMedSeg with MedCLIPSeg within this regime reveals two distinct patterns. On BrainMRI and SkinDerm, UniMedSeg increases OOD-DSC by 1.55–12.87 pp and 8.80–15.10 pp, respectively, while ID-DSC varies by less than 0.4 pp. This results in HM gains of 2.08–16.10 pp and 9.11–16.68 pp, corresponding to the near-vertical arrows toward higher OOD-DSC in Figure 3(b). This behavior aligns with the role of cross-layer uncertainty propagation, which suppresses unreliable vision–language alignment under distribution shift; ablation studies confirm that removing this mechanism reduces OOD-DSC by 1.18 pp (Section 4.5). Conversely, on BreastUS and PolypEndo, UniMedSeg yields marginal ID-DSC improvements (+0.01 to +0.79 pp) but degrades OOD-DSC, causing HM drops of 2.55–5.32 pp and 0.65–9.80 pp, respectively. Overall, while UniMedSeg’s reliability modeling enhances cross-modal transfer for BrainMRI and SkinDerm specialists, a single training modality is insufficient to guarantee such benefits, as evidenced by the BreastUS and PolypEndo results. This underscores the necessity of exposing the model to diverse modalities during training—as implemented in Unified-15DS—to achieve consistent performance gains across all domains.

#### 4.4.2. Unified Training Settings

These settings assess generalization to an unseen dataset and joint fitting of heterogeneous data, respectively. In Unified-15DS, EUS is held out for evaluation. All six operating points exceed the HM = 70 contour in Figure 3(b), with HM values of 70.39–81.53%. Although EUS is unseen during training, its ultrasound modality is represented in the training set, indicating that UniMedSeg transfers shared representations across datasets and organs within a known modality.

At 25% of the training data, UniMedSeg achieves an HM of 70.39% and an OOD-DSC of 60.08%, respectively 6.95 and 10.25 pp below MedCLIPSeg (77.34% and 70.33%). This suggests that the proposed reliability and frequency priors require sufficient supervision to benefit generalization across heterogeneous datasets. With 50% and 100% of the data, however, UniMedSeg surpasses MedCLIPSeg by 7.21 and 1.68 pp in HM, with OOD-DSC gains of 11.76 and 2.82 pp, while ID-DSC changes by only +0.01 and +0.07 pp, confirming that cross-dataset generalization improves without compromising in-distribution accuracy. The best HM reaches 81.53% at 100% data.

In Unified-16DS, all 16 datasets are used for training and evaluated in-distribution, as shown in Figure 3(c). UniMedSeg improves ID-DSC over MedCLIPSeg by 0.39 and 0.55 pp at 25% and 50% data, respectively, and is only 0.28 pp lower at 100% data (88.20% vs. 88.48%). Overall, UniMedSeg maintains competitive in-distribution accuracy while providing stronger generalization under unified multi-modal training.

### 4.5. Ablation Study

Table 4 reports leave-one-component-out ablations in the 6DS multi-source setting. Each variant is trained with three random seeds and evaluated on six source (ID) and ten target (OOD) datasets.^1^

**Table 4.** Leave-one-out ablation on 6DS (DSC, %; mean*±* std over three seeds). Sig.: two-sided Wilcoxon signed-rank test against the full model on seed-averaged per-sample DSC. ^∗∗∗^ *p<*0.001; ^∗∗^ *p<*0.01; ns: not significant (*p*≥0.05).

| Variant | ID | OOD | HM | Sig. (ID/OOD) |
| --- | --- | --- | --- | --- |
| **Full UniMedSeg (Ours)** | **88.71±0.30** | **80.97±0.58** | **84.67±0.46** | — |
| w/o Modality Embedding | 88.25±0.46 | 80.65±0.56 | 84.28±0.51 | *** / ns |
| w/o Uncertainty Propagation | 88.25±0.79 | 79.79±1.35 | 83.80±1.04 | *** / *** |
| w/o Freq.-Spatial Dual-Prompt | 88.40±0.21 | 80.29±1.59 | 84.14±0.92 | *** / *** |
| w/o Boundary Contrastive | 88.30±0.24 | 79.47±0.24 | 83.66±0.19 | ns / *** |
| Decoder → MedCLIPSeg | 88.53±0.21 | 80.42±0.75 | 84.28±0.47 | ** / *** |

1. **Component-wise contributions**. Removing any component reduces ID-DSC, OOD-DSC, and HM. The largest HM reductions occur without boundary contrastive sharpening and cross-layer uncertainty propagation (1.01 and 0.87 pp, respectively), followed by frequency-spatial dual-prompt decoding (0.53 pp), the modality embedding (0.39 pp), and replacement of the decoder with that of MedCLIPSeg (0.39 pp). Eight of the ten ID/OOD comparisons are statistically significant, confirming that each component contributes to the final performance.
2. **Generalization effects**. Except for the modality embedding, every ablation causes a larger OOD than ID reduction. Removing boundary contrastive sharpening decreases ID/OOD-DSC by 0.41/1.50 pp, and removing cross-layer uncertainty propagation by 0.46/1.18 pp. The former significantly affects OOD-DSC but not ID-DSC (*p* ≥ 0.05), supporting its role in improving transferability rather than in-domain fitting. Removing frequency-spatial dual-prompt decoding reduces ID/OOD-DSC by 0.31/0.68 pp, and decoder replacement by 0.18/0.55 pp. These results indicate that cross-layer uncertainty propagation, frequency-spatial dual-prompt decoding, and boundary contrastive sharpening primarily improve cross-domain generalization.
3. **Modality adaptation and stability**. Removing the modality embedding reduces ID-DSC by 0.46 pp but OOD-DSC by only 0.32 pp, the latter change not being statistically significant. This pattern is consistent with its role in preserving modality-specific representations for in-distribution segmentation. Removing cross-layer uncertainty propagation or frequency-spatial dual-prompt decoding increases the across-seed standard deviation of OOD-DSC from 0.58 to 1.35 and 1.59, respectively. Overall, modality embedding primarily supports in-distribution adaptation, whereas uncertainty propagation, frequency-spatial decoding, and boundary contrastive learning provide the main gains in generalization and stability.

### 4.6. Qualitative Results

Figure 4 compares UniMedSeg with MedCLIPSeg on one test image per modality. Both models are trained under the identical Unified-16DS protocol (100% data, seed 666), so differences arise solely from the architecture. As the two models attain similar average ID-DSC in this setting (88.20 vs. 88.48%; Figure 3(c)), we show representative cases with moderate per-image gains (2.7–6.1 pp) to illustrate typical error patterns rather than extreme differences.

**Figure 4:**
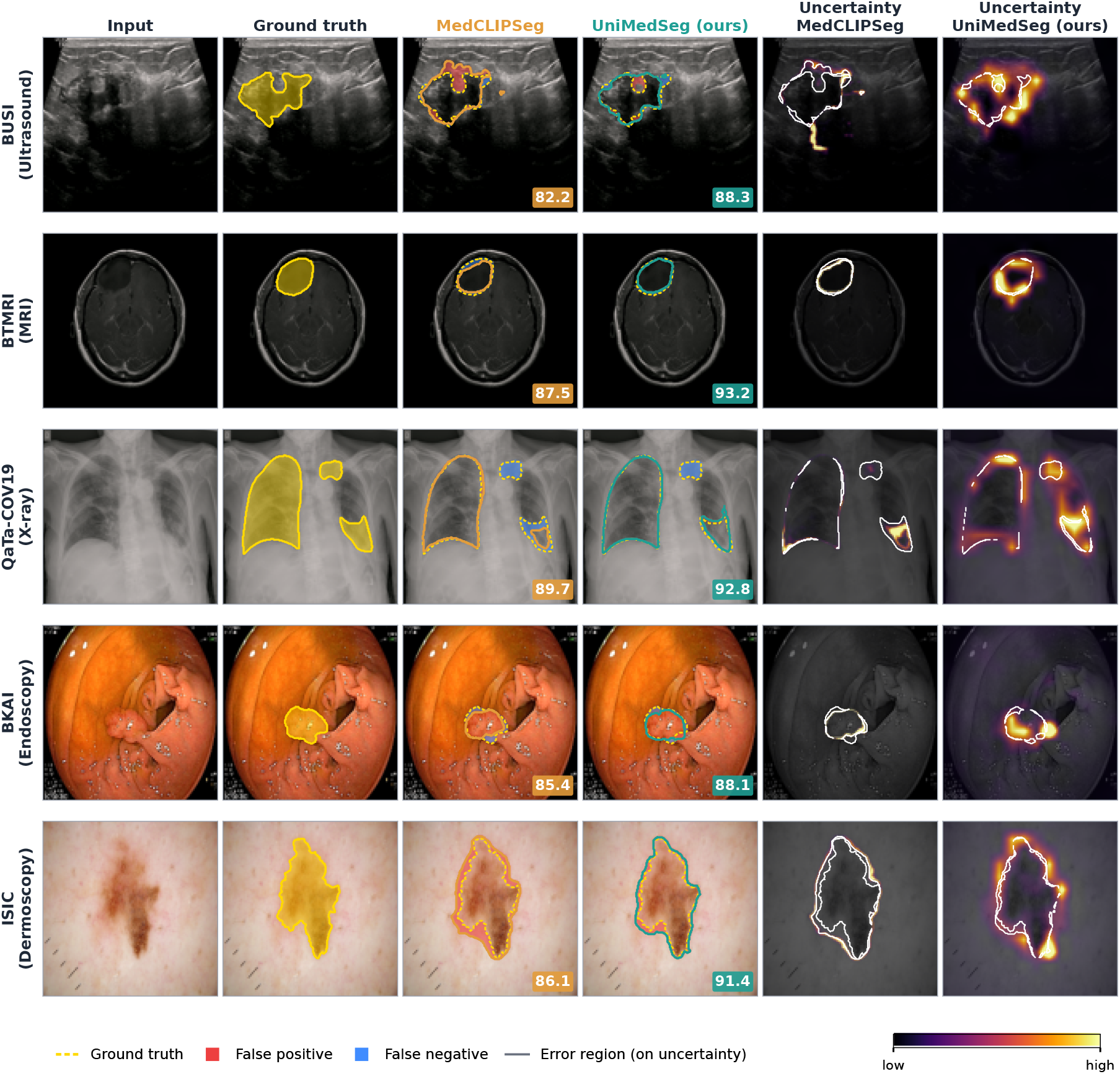
Qualitative comparison of MedCLIPSeg and UniMedSeg, both trained under the Unified-16DS protocol (100% data, seed 666), on representative test images from five modalities. Columns 3–4: predictions (solid contour) against the ground truth (dashed yellow), with false positives in red and false negatives in blue; numbers give the per-image DSC (%). Columns 5–6: uncertainty maps with the segmentation error region outlined in white. MedCLIPSeg uncertainty is the pixel-wise standard deviation over 30 stochastic forward passes; UniMedSeg uncertainty is the fused map *U*. Each map is normalized per image, so only spatial patterns are comparable across methods.

1. **Segmentation Analysis**. The errors of Med-CLIPSeg (Figure 4, column 3) follow two patterns. First, it over-extends into adjacent tissue of similar appearance: in BUSI, a false-positive region (red) extends above the lesion into the surrounding hypoe-choic tissue and an isolated spurious blob appears to its right; in ISIC, a false-positive band surrounds almost the entire lesion boundary on the pigmented skin. Second, it under-segments low-contrast targets: in QaTa-COV19 it recovers only 38% of the lower-right opacity (image coordinates), and in BKAI and BTMRI it misses thin rims along the lesion boundary (blue). UniMedSeg (column 4) substantially reduces both error types, lowering the false-positive area by 43–61% in BUSI and ISIC and recovering 82% of the lower-right lung opacity. The tighter contours are consistent with boundary contrastive sharpening, which separates boundary-foreground from boundary-background representations, and with the separate structure and detail pathways of frequency-spatial dual-prompt decoding. A failure case common to both models is the small upper-right opacity in QaTa-COV19, which neither model detects.
2. **Uncertainty Analysis**. The uncertainty of MedCLIPSeg (column 5) is concentrated in thin bands along its own predicted contour and leaves most of its error regions (white outlines) unmarked, e.g., the false-positive band in ISIC and the upper false-positive region in BUSI. In contrast, the uncertainty map *U* of UniMedSeg (column 6) extends over the regions where errors actually occur, such as the upper lobe of the breast lesion, the lower-right lung opacity, the bottom rim of the polyp, and the upper and lower tips of the skin lesion. Quantitatively, with a normalized-uncertainty threshold of 0.3, the uncertainty of UniMedSeg covers 81–100% of its error pixels across the five cases, compared with 20–51% for MedCLIPSeg, although the pixel-level ranking of errors (AUROC 0.91–0.99) is comparable for both. Most notably, UniMedSeg assigns high uncertainty to this missed upper-right opacity (mean normalized uncertainty 0.69 vs. 0.10 for MedCLIPSeg), flagging a lesion that the segmentation itself fails to capture. This behavior reflects the design of cross-layer uncertainty propagation and precision-weighted fusion trained with the calibration objective *L*_cal_, and indicates that the uncertainty map can direct clinical review to regions that require verification.

## 5. Discussion

The results show that unified multi-modal training can retain source-domain accuracy while improving cross-dataset generalization. In the 6DS multi-source setting, UniMedSeg matches specialist ID-DSC (89.35 vs. 89.55%) and achieves comparable OOD-DSC (81.43 vs. 81.06%), which we attribute to cross-domain knowledge transfer enabled by modality conditioning — the model learns shared representations across datasets while maintaining modality-specific adaptation through the learnable embedding. In the Unified-15DS setting, the best ID–OOD harmonic mean reaches 81.53%, against a maximum of 57.17% for domain-specific specialists tested on unseen modalities. The gap is large in part because the two comparisons involve different OOD tasks: specialist OOD spans multiple modalities, whereas Unified-15DS holds out a single dataset from a seen modality. Nonetheless, it illustrates that joint training over heterogeneous data can substantially reduce the dependency on modality-specific priors.

Ablation results show that the proposed components serve complementary functions. The modality embedding mainly strengthens modality-specific in-distribution representations, while cross-layer uncertainty propagation continuously refines vision–language alignment reliability across encoder layers. Frequency-spatial dual-prompt decoding separately models global structure and fine boundary detail, and boundary contrastive sharpening raises discriminability in ambiguous transition regions. The disproportionate OOD reductions when removing uncertainty propagation (−1.18 pp OOD vs. −0.46 pp ID) and boundary contrastive sharpening (−1.50 pp OOD vs. −0.41 pp ID, with the ID change not statistically significant) confirm that these components primarily benefit generalization rather than in-domain fitting. The increased across-seed variance in OOD-DSC without uncertainty propagation or frequency-spatial decoding (0.58 to 1.35 and 1.59) further suggests that explicit reliability and frequency modeling reduce sensitivity to initialization. Unified training is not uniformly preferable. On ColonDB and BKAI, the unified model falls below the specialist by 2.03 and 4.77 pp, and at 25% training data the Unified-15DS UniMedSeg trails MedCLIPSeg in OOD-DSC. Datasets with distinctive acquisition statistics and low-data regimes may therefore still benefit from domain-specific fine-tuning or modality-adaptive capacity.

One limitation remains open: the frequency cut-off ratio and the structure–detail fusion gate are shared across modalities, and the experiments use two-dimensional data only; modality-adaptive frequency modeling and extension to three-dimensional volumetric segmentation are natural next steps.

## 6. Conclusion

We presented UniMedSeg, a unified multi-modal medical image segmentation framework that combines two core components—cross-layer uncertainty propagation for continuously refining vision–language alignment reliability, and frequency-spatial dual-prompt decoding for separately modeling global structure and fine boundary detail—together with a boundary contrastive loss that improves discriminability in ambiguous transition regions. Experiments on 16 datasets spanning five modalities demonstrate that UniMedSeg retains source-domain accuracy comparable to domain-specific models while substantially improving cross-dataset generalization in the Unified-15DS setting (best HM-DSC 81.53% versus 57.17% for domain-specific models tested on unseen modalities). These results suggest that combining modality conditioning, cross-layer reliability propagation, and structure–detail decoupling provides a practical path toward a single model that achieves competitive segmentation performance, cross-dataset generalization, and spatially calibrated uncertainty.

## CRediT authorship contribution statement

**Jingling Zhang:** Conceptualization, Methodology, Software, Investigation, Writing - Original Draft, Visualization. **Shuting Zheng:** Software, Validation, Data Curation, Visualization. **Xiangfei Liu:** Methodology, Formal Analysis, Validation. **Wen Zhang:** Writing - Review & Editing, Supervision, Resources. **Jia Gu:** Conceptualization, Supervision, Project Administration, Funding Acquisition.

## Data availability

The datasets used in this study are publicly available. The code and models are openly accessible at https://github.com/699zjl/UniMedSeg.

## Acknowledgments

This work was supported by the Science and Technology Development Fund of Macao under Grant No. 0002/2024/RIA1, and by Fujian Province Young and Middle-aged Teachers’ Educational Science Research Projects (Science and Technology Category) under Grant No. JAT232014.

---

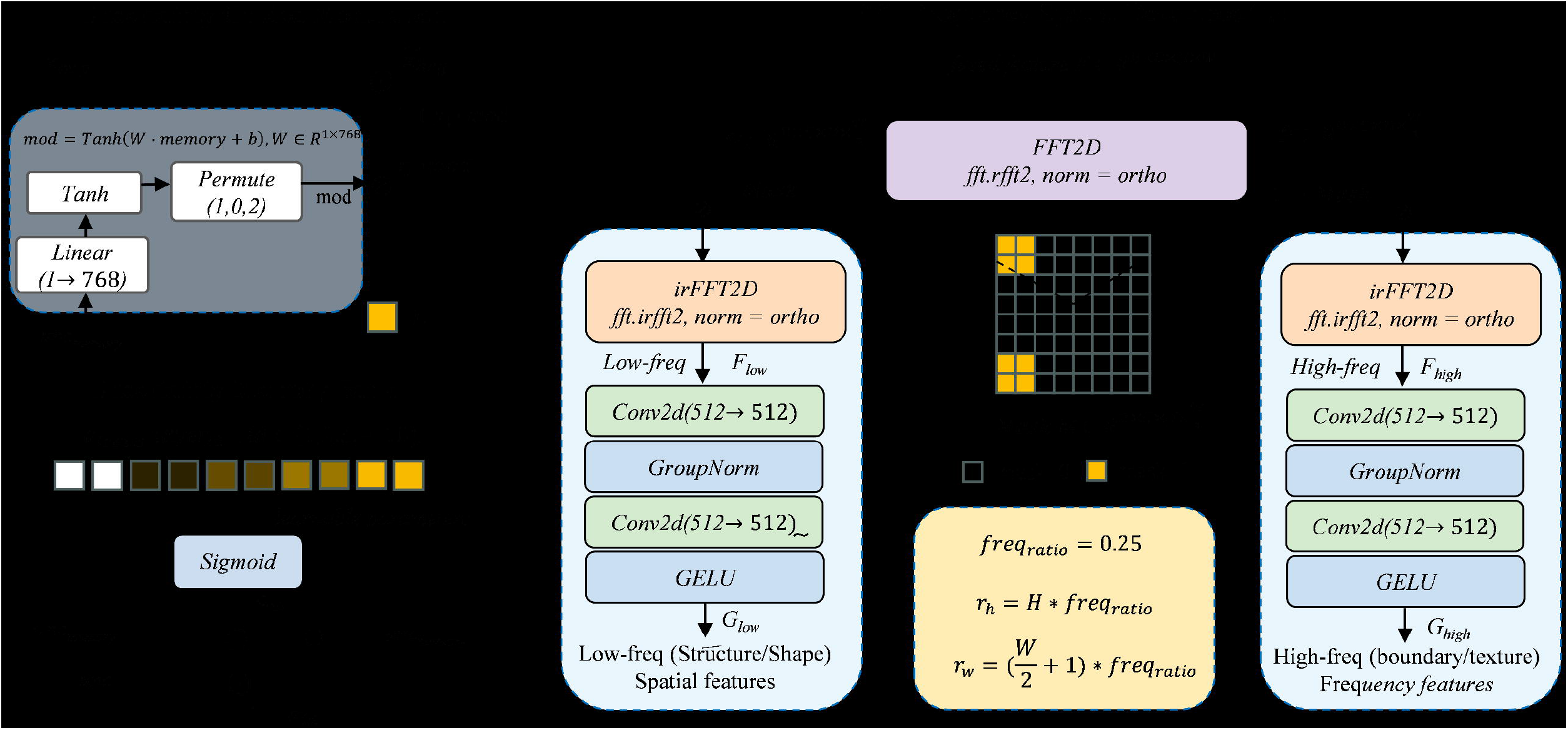

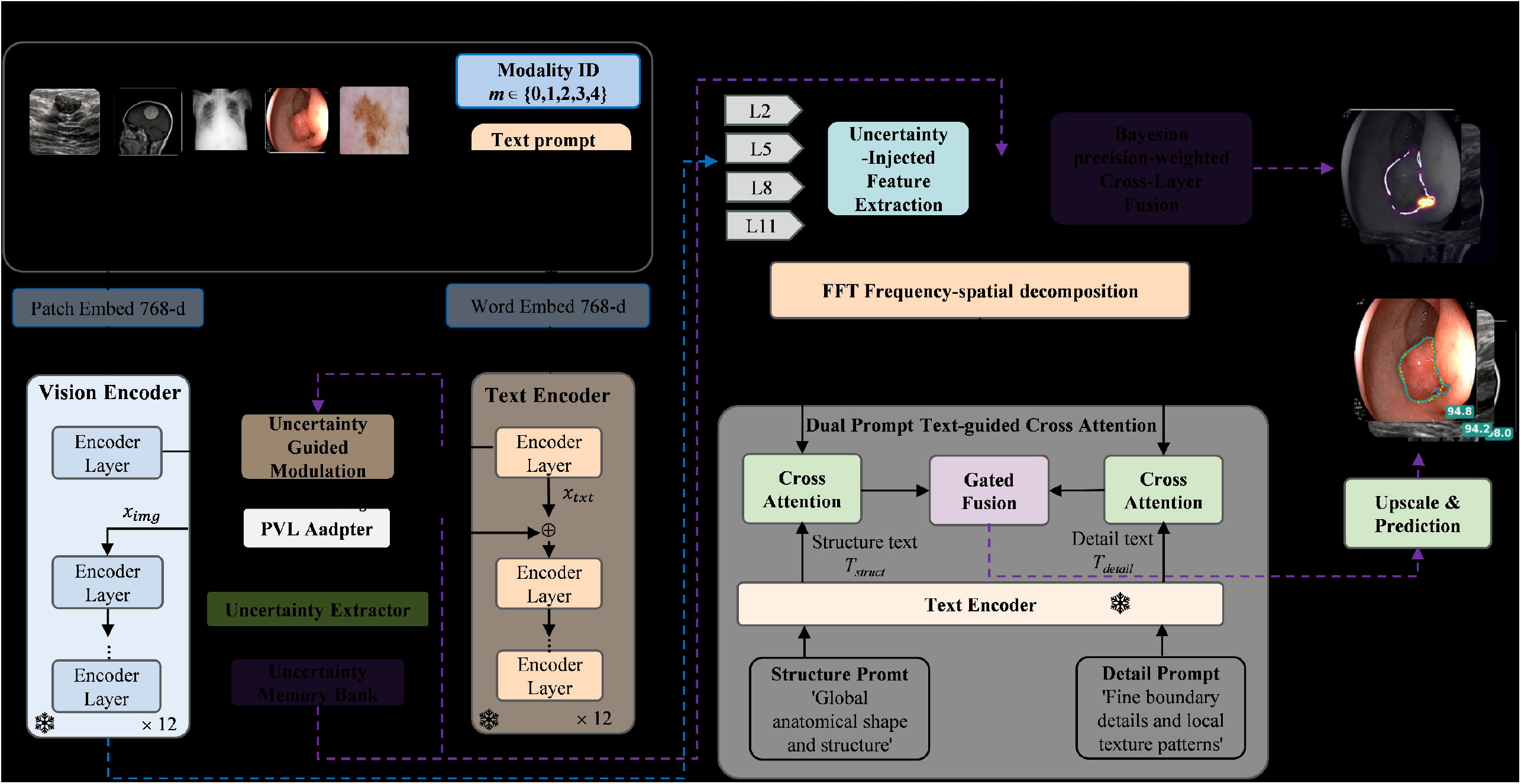

## Footnotes

1 Tables 2 and 3 report a single run with seed 666, whereas the present evaluation uses three seeds and includes QaTa-COV19 and EUS in the ID average.

